# Influence of functional and phylogenetic dissimilarity on exotic plant invasion depends on spatial scale

**DOI:** 10.1101/2023.08.18.553820

**Authors:** Maria A. Perez-Navarro, Adam T. Clark, Joshua I. Brian, Harry E. R. Shepherd, Jane A. Catford

## Abstract

1. Darwin’s theory of natural selection provides two seemingly contradictory hypotheses for explaining the success of introduced species: 1) the pre-adaptation hypothesis posits that introduced species that are closely related to native species will be more likely to succeed than distantly related invaders because they already possess relevant characteristics; 2) the limiting-similarity hypothesis posits that invaders that are more similar to resident species will be less likely to succeed due to competitive exclusion. Previous studies assessing this conundrum show mixed results, possibly stemming from variation in study spatial scales and lack of both functional and phylogenetic information.
2. We used species abundances compiled in a 33-year grassland successional survey based at Cedar Creek Ecosystem Science Reserve (USA) to assess the support for the pre-adaptation and limiting similarity hypotheses at two different spatial scales (neighbourhood scale of 0.5m^2^, site scale of ~40m^2^). We combined compositional surveys of 303 vascular plant taxa (256 native, 47 introduced) taken across 2700 plots in a chronosequence of abandonment from agriculture with species functional dissimilarities, phylogenetic distances, environmental covariates and information on species origin.
3. Our results consistently supported the pre-adaptation hypothesis at the site scale but diverged at neighbourhood scale, with functional dissimilarity supporting the limiting similarity hypothesis and phylogenetic distance supporting the pre-adaptation hypothesis. Introduced species with low leaf dry matter content (LDMC), low height and high seed mass tended to be most abundant than rest of species, while relationships between species abundance and specific leaf area (SLA) varied with scale. Introduced species were more abundant than natives at higher concentrations of soil N but were less abundant than natives over time.
4. *Synthesis:* Our study highlights the importance of environmental filtering on grassland community assembly at the scale of a site – here 40 m^2^, a spatial resolution that is usually considered “local”. This influence of environmental filtering might mask effects of limiting similarity at small “local” scales. Our results demonstrate the importance of accounting for both phylogenetic and functional dissimilarity when examining the complex interaction between species biogeographic origin, functional strategies and evolutionary history as considering one alone can lead to different conclusions.

## Introduction

Humans are introducing species beyond their native ranges at an ever-increasing rate (Seebens et al., 2017) with a consequent rise in the impact of invasions on biodiversity, ecosystem function, human health, and economy (Mazza, 2018; Pejchar & Mooney, 2009; Pyšek, Hulme, et al., 2020). Despite considerable research into the characteristics and processes underlying the establishment and spread of introduced (exotic, non-native, alien) species, much uncertainty about species invasion remains (Catford et al., 2019; Pyšek, Novoa, et al., 2020; Pyšek & Richardson, 2008). From the wide number of hypotheses attempting to explain species invasions (Catford et al., 2009), dissimilarity between introduced and native species and associated hypotheses about pre-adaptation and limiting similarity are key (Catford et al., 2009; Enders et al., 2020; Lemoine et al., 2016).

The pre-adaptation and limiting similarity hypotheses are deeply rooted in Natural Selection Theory and are known as Darwin’s naturalization conundrum, owing to their contrasting expectations regarding functional and phylogenetic similarity and invasion success (Cadotte et al., 2018; Darwin, 1859; Diez et al., 2008; Thuiller et al., 2010). The pre-adaptation hypothesis is predicated on the importance of environmental filtering for community assembly. It predicts that introduced species more closely related to native species will be more successful than distantly related invaders because they are more likely to already possess traits that are suitable in the introduced range (Fridley & Sax, 2014; Lososová et al., 2015; Qian & Sandel, 2022). In contrast, the limiting similarity hypothesis (also known as Darwin’s naturalization hypothesis) supposes that biotic interactions, particularly competition, dominate community assembly. As such, more distantly related and functionally dissimilar introduced species will be more successful than closely related ones as they will be better able to avoid competition with natives and fill “empty niches” (Carboni et al., 2013; Catford et al., 2009; Fridley & Sax, 2014; Thuiller et al., 2010). These hypotheses are not mutually exclusive, but support for them has been mixed and inconsistent (Cadotte et al., 2018; Gallien & Carboni, 2017).

Evidence for the pre-adaptation and limiting similarity hypotheses may vary across spatial scales reflecting the differential influence of environmental filtering and competition on community assembly (Cadotte et al., 2018; Gallien & Carboni, 2017; Götzenberger et al., 2012; Park et al., 2020). Generally, higher support for limiting similarity is expected at finer spatial resolutions, as greater environmental homogeneity at these scales increases our ability to detect direct plant-plant interactions (Fig. 1) (Cadotte et al., 2018; Kraft et al., 2015). In contrast, greater support for the pre-adaptation hypothesis is expected at larger spatial scales, as increased environmental heterogeneity increases the strength of environmental filtering relative to biotic interactions (Fig. 1) (Thuiller et al., 2010). However, a recent study revealed inconsistent patterns with this expectation (Cadotte et al., 2018). Among other reasons, the observed lack of consistency could be due to the way spatial scales are described, grouped and subsequently compared, leading to a form of apparent context dependence (*sensu* Catford et al., 2022). For example, scales varying from 1 m^2^ to over 100 m^2^ are frequently described as “local” (Cadotte et al., 2018; Cadotte, Borer, et al., 2010; Cadotte, Cavender-Bares, et al., 2009; Marx et al., 2016), thus likely capturing different ecological processes within the same spatial category.

**Figure 1.**
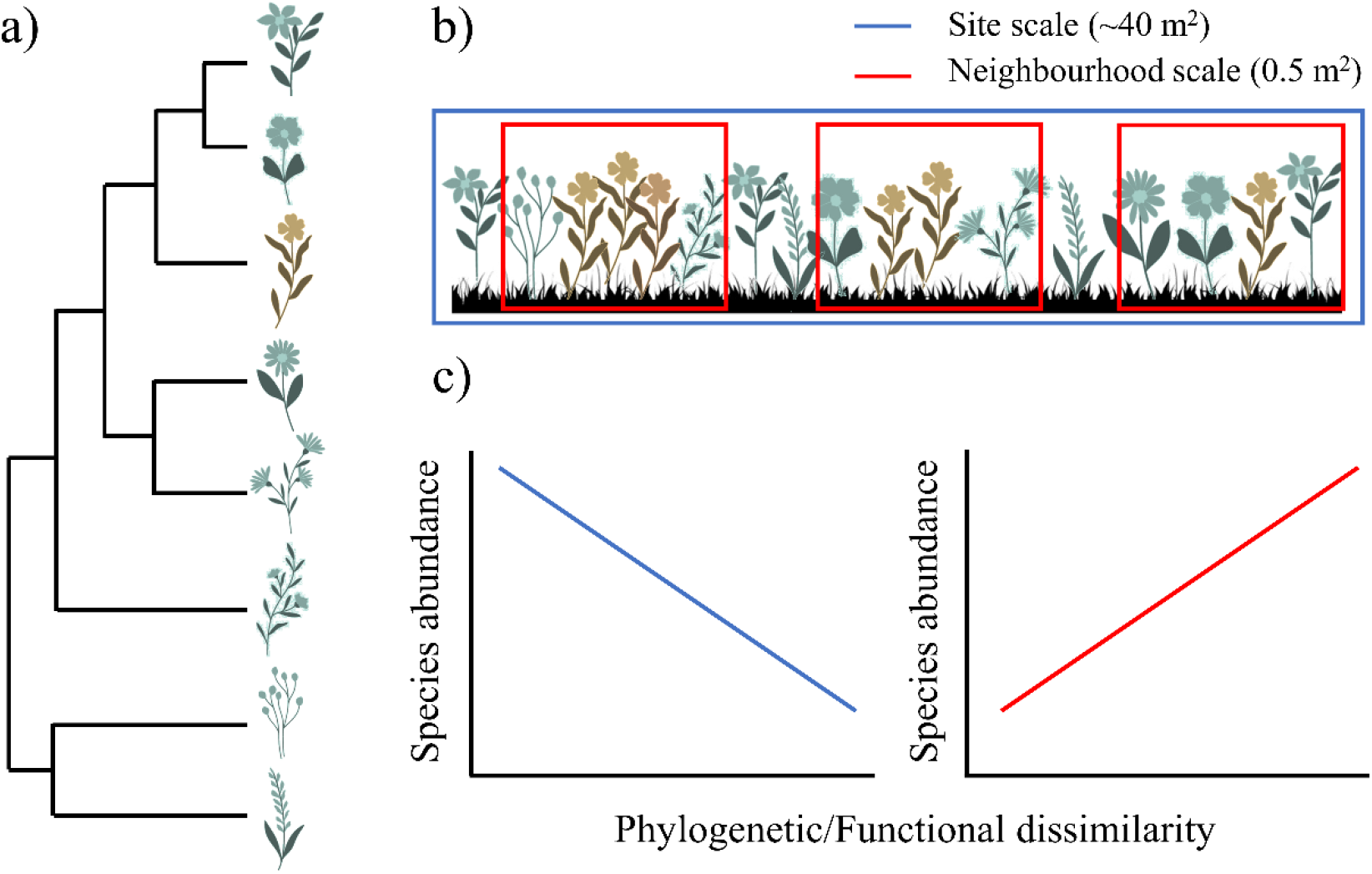
Summary diagram of expected support for pre-adaptation and limiting similarity hypotheses at different spatial scales (i.e., spatial resolutions or grains). Each plant icon represents a different hypothetical herb species. Dark green colour represents native species while ochre represents introduced species. Panel “a” shows the phylogenetic distribution of the species. Panel “b” represents the spatial distribution of the species in a given transect. Dark blue box represents the site scale (defined in this study as 40 m^2^). Red boxes represent the neighbourhood scale (defined in this study as 0.5 m^2^). Panel “c” represent the expected relationship between species abundances and their phylogenetic/functional dissimilarity to the rest of the species present in the community. Dark blue trend at site scale would support the pre-adaptation hypothesis (also known as environmental filtering) and red trend at neighbourhood scale would support the limiting similarity hypothesis (also known as Darwin’s naturalization hypothesis).

Although it is assumed that individuals only directly interact with close neighbours (Götzenberger et al., 2012; Stoll & Weiner, 2000), there is no defined spatial resolution (or grain, *sensu* Wiens, 1989) that delimits the spatial scale at which these interactions occur, as it depends on the size of the organisms and micro-habitat conditions. For herbaceous plants, it has been suggested that the “neighbourhood scale” ranges from a few centimetres to around one square metre (Götzenberger et al., 2012). Beyond this scale, the relative importance of environmental filtering (i.e. pre-adaptation) is expected to increase, reducing the ability to detect effects of biotic interactions (i.e. limiting similarity) (Gotelli & Mccabe, 2002; Watkins et al., 2003). Nonetheless in most studies of plant invasions, smaller spatial scales range from 1 to 10s of meters, while regional scales typically comprise extents from a few km to continental scales (Cadotte et al., 2018; Cadotte, Hamilton, et al., 2009; Lososová et al., 2015). This means that, for herbaceous plants, there are very few studies that truly examine relationships at the neighbourhood scale (i.e., less than 1 m^2^), where limiting similarity is more likely to be detected. A robust test of Darwin’s naturalization conundrum, that clearly disentangles the influence of competition and environmental filtering on invasion success, requires analysis of finer spatial resolutions.

Multiple studies have considered phylogenetic dissimilarity to explain invasion success, but studies that combine phylogenetic data with multivariate and univariate functional data are rare (Cadotte et al., 2013). Although closely related species are generally expected to have similar functional traits, functional convergence and divergence can decouple this relationship, highlighting the need to consider both functional and phylogenetic information when assessing Darwin’s naturalization conundrum (Cadotte et al., 2017). The use of ranked functional dissimilarities, which emphasize the invader’s hierarchical position on each trait gradient, could also indicate when functional differences are associated with higher competitive capacities (Carboni et al., 2016; Gallien et al., 2015).

Co-variation in other factors, such as environmental conditions and historical land use, may also contribute to apparent inconsistencies between studies examining Darwin’s naturalization conundrum (Gallien & Carboni, 2017; Ma et al., 2016). For example, low resource availability might act as an environmental filter on community assembly (Aerts, 1999; Conradi et al., 2017), strengthening support for the pre-adaptation hypothesis. Alternatively, past disturbances, such as changes in land use or fire, can open a window of opportunity for the colonization of new species, regardless of their functional similarity to natives (Catford et al., 2012; Kneitel & Perrault, 2006).

Most studies addressing Darwin’s naturalization conundrum have only examined it in relation to introduced species, disregarding whether the conundrum also applies to co-occurring native species (Cadotte et al., 2018; Ma et al., 2016). Considering both native and introduced species can reveal whether all colonising species follow the same set of assembly rules and life history strategies regardless of their species origin, or whether some trends are applicable to introduced species only (Lemoine et al., 2016). If the two groups of species do follow the same set of assembly rules, then the conundrum that has been focused on non-native naturalization is in fact universal for plant colonization, and the implications are broader than simply predicting the establishment of non-native species in new communities.

Working in secondary successional grasslands at Cedar Creek Ecosystem Science Reserve, Minnesota, USA, here we use a long-term survey (1983-2016) to determine support for the two hypotheses associated with Darwin’s naturalization conundrum – the pre-adaptation and limiting similarity hypotheses. We work at spatial scales that correspond with the underlying competition and environmental filtering processes, using 0.5 m^2^ plots to represent the neighbourhood scale and ~40 m^2^ transects to represent the site scale. To test the two competing hypotheses, we examine relationships between functional and phylogenetic dissimilarities and native and introduced species’ abundances at the two spatial resolutions. We have two expectations:

1. the limiting similarity hypotheses will gain most support at the neighbourhood scale (0.5 m^2^) where individuals can directly interact and compete for resources, whereas the pre-adaptation hypothesis will show higher support at site scale (~40 m^2^) where environmental filtering is likely to be stronger (Fig. 1); and
2. introduced and native species will show similar responses to functional, phylogenetic and environmental predictors, though introduced species might have different life history traits and higher competitive abilities than native species based on potential evolutionary differences (Fridley & Sax, 2014; Gallien et al., 2015; Zheng et al., 2015).

Our approach tries to clarify several uncertainties that are often overlooked in other studies examining Darwin’s naturalization conundrum. We do this by: i) examining trends at unusually fine spatial resolutions; ii) using an unusually comprehensive analytical approach that considers phylogenetic, multivariate and univariate functional indices and environmental covariates; and iii) investigating support for the pre-adaptation and limiting similarity hypotheses in native species as well as introduced species.

## Materials and Methods

### Study system

The study is based on data collected in a long-term vegetation survey carried out at Cedar Creek Ecosystem Science Reserve, Minnesota (hereon Cedar Creek; 45.4° N, 93.2° W), within the experiment E-014 (https://www.cedarcreek.umn.edu/research/data/methods?e014). The study site is characterized by nitrogen-limited sandy soils, annual precipitation of 775 (±158 *SD*) mm, mainly concentrated between April and August, mean summer temperatures of 27 °C (±1.6 SD), and winter lows of −14 (±1.2 SD) °C (according to closest weather station for the reference climate period 1963 to 2016), (see Inouye et al., 1987 for further details on site history).

Before widespread land clearing in the mid-19th century, the study region was composed of a mix of grassland, oak savanna, deciduous forests, and wetlands (Cushing, 1963). Current vegetation in experiment E-014 is characterized by grasslands, with dominance of species from families *Asteraceae, Brassicaceae, Cyperaceae, Fabaceae, Poaceae, Polygonaceae,* and *Rosaceae*. Both native and introduced species are present, with introduced species contributing an average of 44% of plant cover across sample years.

### Field sampling and data preparation

Experiment E-014 is divided into 26 fields, which were previously used as cropland for various species (including soybeans, rye, oats and corn) and were abandoned in a staggered sequence between 1927 and 2015. The exact year of abandonment was obtained from historical documents and aerial photographs. Within each field, 4 permanent 39 m-long transects (hereafter site scale) were laid out, except for 2 fields where 6 transects were established. Each transect was divided into 25 plots of 0.5 m x 1 m (hereafter neighbourhood scale), located every 1.5 m along the transect (with the first plot located at 1.5 m from the beginning of the transect) (Fig. S1). Heavy afforestation has occurred in all survey plots of one field and partially in plots of three other fields. Trees were absent or rarely appeared in the remaining 22 fields. When trees appeared, they were discarded from the dataset, to keep only grassland communities. Finally, in all but four fields, half of the plots in each field were exposed to experimental periodic burning treatments since 2006, with a frequency of approximately one fire every three years (Clark et al., 2018).

Within the 0.5 m^2^ plots, percent cover of every plant species, bare ground and litter were visually estimated (see Clark et al., 2018 for further details). Eight vegetation surveys were conducted between 1983 and 2016, with a survey frequency of approximately 6 years (Clark et al., 2018). In total, 2700 plots (distributed within 108 transect, nested within 26 fields) were sampled, although some plots were not sampled in every survey. Across all years a total of 16,944 plots surveys were conducted, encompassing 303 vascular plant species from which 218 were native, 47 were introduced and 38 had unknown origin (according to the species origin status recorded by the Agriculture Department of United States; USDA-NCRS, 2022). This last category included both species with unknown origin status and species identified at genus level, the genera of which contain both introduced and native species in Minnesota.

For each plot, soil nitrogen, soil carbon, soil organic matter concentration and light penetration were measured. Though these data were sampled in multiple years, we used average information across time, as soil and light sampled years do not match plant abundance surveys. In addition, we also estimated each species’ minimum colonization time in each community. minimum colonization time (hereon colonization time) is similar to the concept of minimum residence time used in invasion ecology (Pyšek et al., 2004; Wilson et al., 2007) but refers to the minimum amount of time a species (whether native or introduced) was present in a given plot following agricultural abandonment. We calculated colonization time as the difference between the year that a species was observed in a given plot and the year when the species first colonised the plot. Finally, succession time (i.e., time since plot abandonment; estimated as the difference between year of plot survey less year of field abandonment) and presence of burning treatment (Yes/No) were also considered in the analyses as a proxies of disturbances (Kneitel & Perrault, 2006).

All described variables were estimated at both neighbourhood and site scales. To obtain data at site scale, we averaged data for percent of plant cover, soil content in carbon, nitrogen and organic matter and light penetration across all plots within the same transect; and re-estimated the minimum colonization time according with transect species composition and time records. Year of abandonment and burning treatment did not require scaling as these were already at the site scale. See Appendix S1 for details of all methods.

### Trait data

Functional trait data for the study species were collected in grasslands and savannahs at Cedar Creek (Cadotte et al., 2009; Catford et al., 2019; Willis et al., 2010 and experiment E-133). We selected seven functional traits with high data coverage across the taxa included in our study: specific leaf area (SLA), plant height, leaf area, leaf dry matter content (LDMC), leaf fresh mass, leaf dry mass (all of them with data for more than 60% of the 303 taxa observed across all study plots and survey years) and seed mass (data for 55% of the observed taxa). These traits cover most of the variability related to plant functional strategies (Díaz et al., 2016). We averaged trait data at species level when more than one individual was sampled per species. Therefore, we only included the 152 taxa for which we had data on all 7 functional traits in our analyses. Although around 50% of the study taxa (i.e., 151 taxa) were not included in functional analyses, trait representativeness at plot level (neighbourhood scale) and transect level (site scale) was high, with most of the plot and transects having trait data for more than 75% of the taxa, both in terms of number of taxa and percentage cover (Fig. S4). See Appendix S2 for details on trait data compilation.

### Functional dissimilarity estimation

We first built a common multivariate functional trait space by using principal component analysis (PCA, package ade4, Dray & Dufour, 2007) to convert the functional space of the 7 selected functional traits into a three-dimensional volume defined by the first three PCA axes. Trait variables were scaled before conducting the PCA. The first three PCA axes explained 89.37% of the total variance (Fig. S5). We selected this multivariate approach since it facilitates an integrated estimate of species functional traits. However, we also estimated ranked univariate functional dissimilarities for each individual trait in recognition of potential effects of specific traits (including how they might relate to competition) and to help with interpretation of the results (Carboni et al., 2016; Gallien & Carboni, 2017).

After conducting the PCA, we obtained three PCA coordinates for each species. With these coordinates we estimated each species dissimilarity to the rest of species in its community by estimating the three-dimensional Euclidean distance to the community functional centroid (equivalent to functional dispersion measurement *sensu* Laliberte & Legendre, 2010). The community functional centroid was estimated as the centre of gravity of the functional axis values of all species present in the community except the target species (i.e., the mean of species’ coordinates in each PCA axes weighted by each species cover percent; Fig. S6).

Similarly, we quantified ranked functional dissimilarity for each species and each individual trait by calculating the difference between each species’ functional trait values and the functional centroid of the rest of the species present in the community. Again, the community centroid was estimated as the mean of the rest of species functional trait values weighted by their percent cover. Therefore, species with positive ranked functional dissimilarity indicates that the species have higher trait values than the average of the rest of the community while those having negative values have lower trait values than the rest of the community (Gallien & Carboni, 2017). Multivariate and univariate functional dissimilarities were estimated both at neighbourhood and site scale according to community composition in each case.

### Phylogenetic distance estimation

We used V.phylomaker package (Jin & Qian, 2019) based on Smith & Brown, 2018 and Zanne et al., 2014 phylogenies to obtain the phylogenetic tree of the total set of species present in the study area (Fig. S7). Species names were harmonised according with The Plant List using the taxonstand R package (Cayuela et al., 2021). From this phylogenetic tree with all the study species, we obtained phylogenetic trees for each plot and transect, and estimated the phylogenetic distance between each species and the rest of the community as the weighted mean of each pairwise phylogenetic distance between the target species and each single species present in the community, weighted by the species’ cover percent (Cadotte, Davies, et al., 2010).

### Statistical analyses

We ran two sets of linear mixed effect models, one at plot level (neighbourhood scale) and another at transect level (site scale) to assess the effect of phylogenetic and functional dissimilarity on species abundance. For both sets of models, we used logarithmically transformed percent cover as the response variable. For explanatory variables, we used multivariate functional dissimilarity, ranked univariate functional dissimilarity (seed mass, SLA, LDMC, leaf area, leaf fresh mass and dry mass and plant height), phylogenetic distances and environmental and historical land use variables (soil nitrogen, soil carbon, soil organic matter content, minimum colonization time, succession time, burning treatment and light penetration) and their interaction with species origin. Species origin was considered as a categorical variable with two categories: “introduced” (including only USDA introduced category) and “native” species (including both native species and species of unknown origin). There were only 16 unknown species from the total of 152 species included in the analyses. Univariate functional trait variables were scaled (subtracting the mean and dividing by standard deviation), and the rest of the variables were logarithmically transformed when required to meet normality of model residuals. From this set of explanatory variables, we built 5 different alternative models for each spatial scale (neighbourhood and site), from model 1 including only two variables to model 4 including all the described variables above (Table S2). In every case, models included at least the phylogenetic distance and functional dissimilarity variables, which were at the core of our analysis of Darwin’s naturalization conundrum hypotheses. In addition, model 5 included the triple interaction of each explanatory variable with species origin and succession time in order to assess the potential effect of succession time on species cover changes (Clark et al., 2018) (Table S2). Collinearity between explanatory variables was tested for each model using Pearson correlation and Variance Inflation Factor analyses (Nakagawa et al., 2022), and highly correlated variables were removed from the fitted models. Species, year and plot nested within transect and transect nested within field were established as crossed random intercepts in case of neighbourhood scale models; and species, year and transect nested within field were established as crossed random intercepts for site scale models. The best model for each spatial scale was selected according to AICc and BIC criteria (Table S3). All analyses and data manipulation were performed using R version 4.2.1 (R core Team, 2022).

## Results

At both site (~40 m^2^) and neighbourhood (0.5 m^2^) spatial scales, the full models, which included phylogenetic distances, multivariate and univariate functional dissimilarities and environmental variables, were the more parsimonious ones according to BIC and AICc indexes (Table S3). Below we summarise the results according to our two expectations outlined in the Introduction. The first section describes findings related to species’ multivariate functional dissimilarity and phylogenetic distance, which we used to test support for the two hypotheses central to Darwin’s naturalization conundrum. The second section compares native and introduced species abundance in relation to ranked univariate functional dissimilarities and environmental variables.

### 1. Support for limiting similarity and pre-adaptation hypotheses across spatial scales

Relationships between species abundance and multivariate functional dissimilarity depended on the spatial scale at which relationships were examined (Figs. 2-5, Tables S5-S6). At the neighbourhood scale, species that were functionally distinct from co-occurring species in their communities reached higher abundances than functionally similar species, similarly for both native and introduced species, consistent with the limiting similarity hypothesis (Fig. 2c). Although not statistically significant for introduced species, species more functionally similar to the rest of the community were more abundant at the site scale (Fig. 2d), consistent with the pre-adaptation hypothesis. In contrast, regarding phylogenetic distance, introduced species that were more closely related to co-occurring species reached higher abundances than distantly related species at both spatial scales, providing support for the pre-adaptation hypothesis (Fig. 2a,b). Native species, however, showed a contrasting trend. The relationship between phylogenetic distance and native species abundance was negligible at the neighbourhood scale, but it was significantly positive at the site scale. As such, native species more distantly related to co-occurring species showed higher abundances than closely related natives (Fig. 2b).

**Figure 2.**
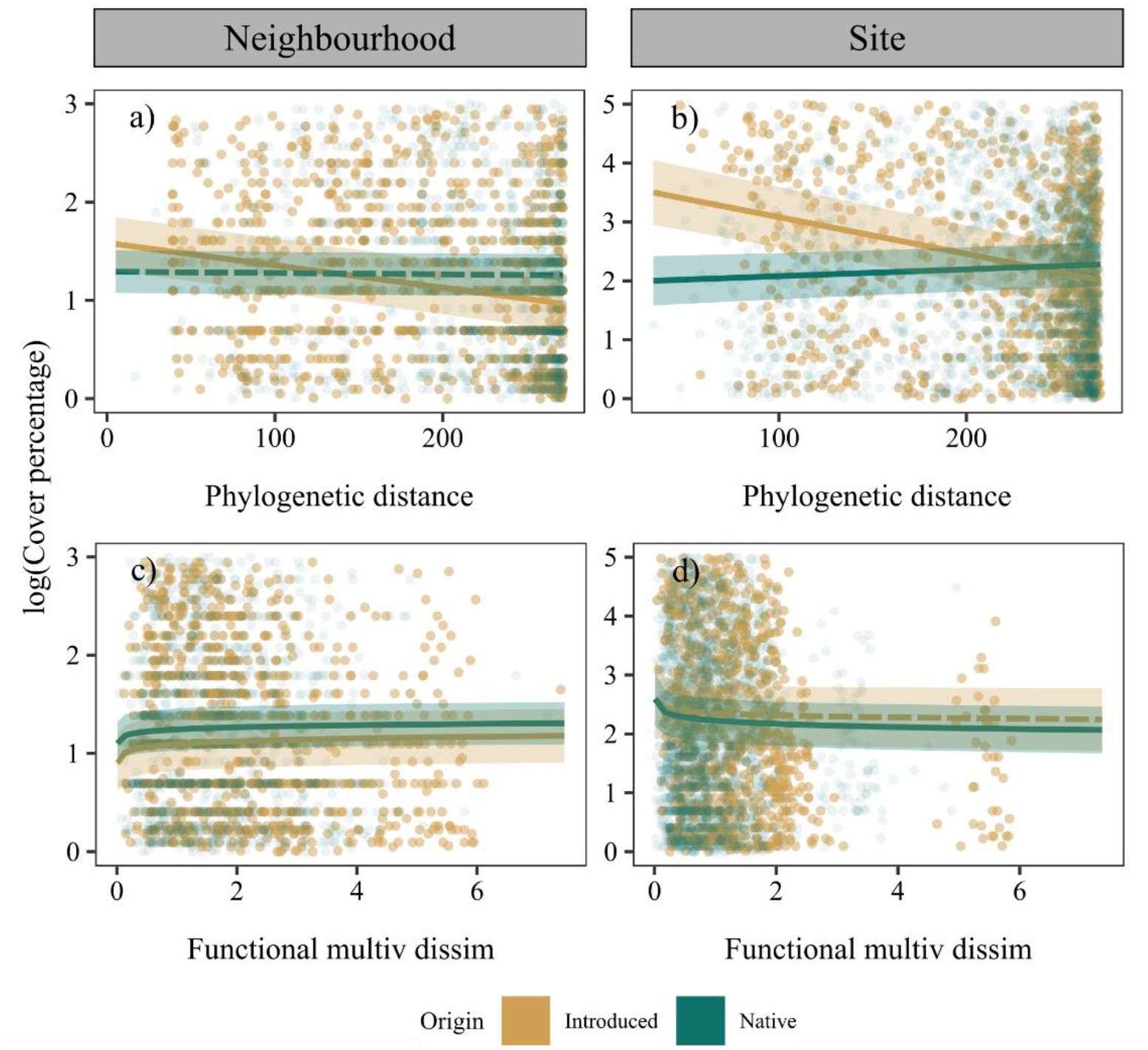
Regression plots from the final selected model for both neighbourhood (0.5 m^2^) and site (~40 m^2^) spatial scales. Y-axes represent log of species cover percent. X-axes correspond to each of the variables used to evince Darwin’s naturalization conundrum (multivariate functional dissimilarity and phylogenetic distance). Dark green colour represents native species populations and ochre colour represents native species populations. Solid and dashed lines represent statistically significant (p < 0.05) and non-significant effects (p > 0.05), respectively. Plots only show 5% of species x plot data used in models at neighbourhood scale and 25% of species x transect data at site scale, in order to reduce the number of points (to 5000 each) and improve visualization.

### 2. Differences between introduced and native species in response to univariate functional traits and environmental conditions

Native and introduced species had distinct relationships with all ranked univariate trait dissimilarities and environmental variables, except for LDMC dissimilarity and soil nitrogen content at the site scale and burning treatment at both the neighbourhood and site scale (Fig. 4). Some of the statistically significant differences between introduced and native species involved differences in relationship sign (direction), but most of these relationships only differed in their strength (magnitude) (Fig. 5).

In the case of ranked trait dissimilarities, introduced and native species had contrasting relationships for LDMC at the neighbourhood scale and seed mass at both scales (Figs. 4-5). While native species showed higher abundances with higher LDMC or lighter seeds, introduced species were more abundant when showing lower LDMC and heavier seeds than rest of species of the community (Fig. 3c-d, Supplementary Material Fig. S8c-d). For the other two ranked trait dissimilarities (height and SLA), introduced and native species’ relationships differed in strength only (Figs. 3a-b & S8a-b). Native and introduced species with shorter heights and higher SLA than the rest of the community tended to be more abundant (Figs. 4-5).

**Figure 3.**
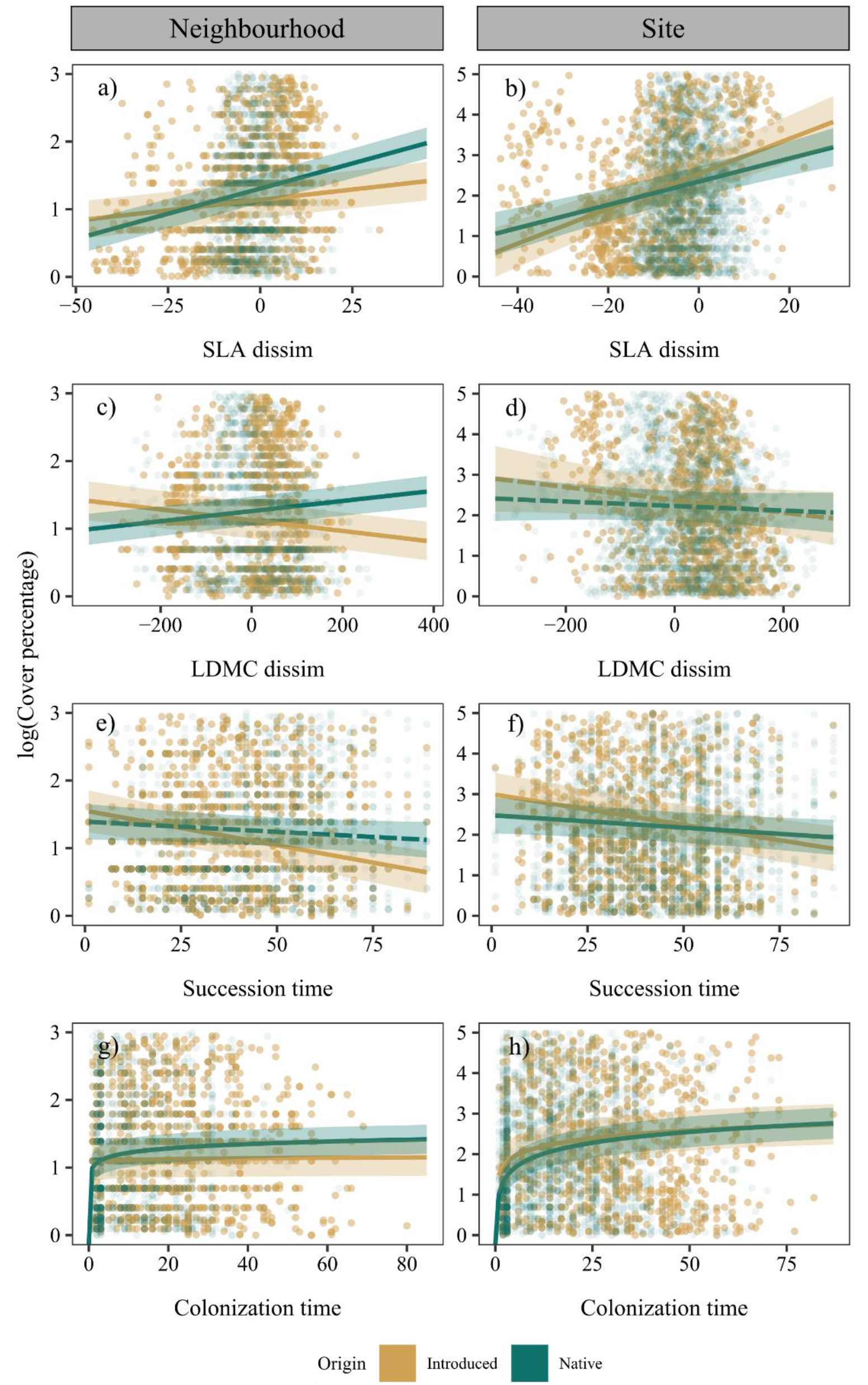
Regression plots from the final selected model for both neighbourhood (5 m^2^) and site spatial scales (~40 m^2^). Y-axes represent log of species cover percent. X-axes correspond SLA and LDMC dissimilarities and succession time and species minimum colonization time. The rest of the traits and explanatory variables included in the final models can be seen in Fig. S8. Dark green colour represents native species populations and ochre colour represents native species populations. Solid and dashed lines represent statistically significant (p < 0.05) and non-significant effects (p > 0.05) respectively. Plots only show 5% of species x plot data used in models at neighbourhood scale and 25% of species x transect data at site scale, in order to reduce the number of points (to 5000 each) and improve visualization.

**Figure 4.**
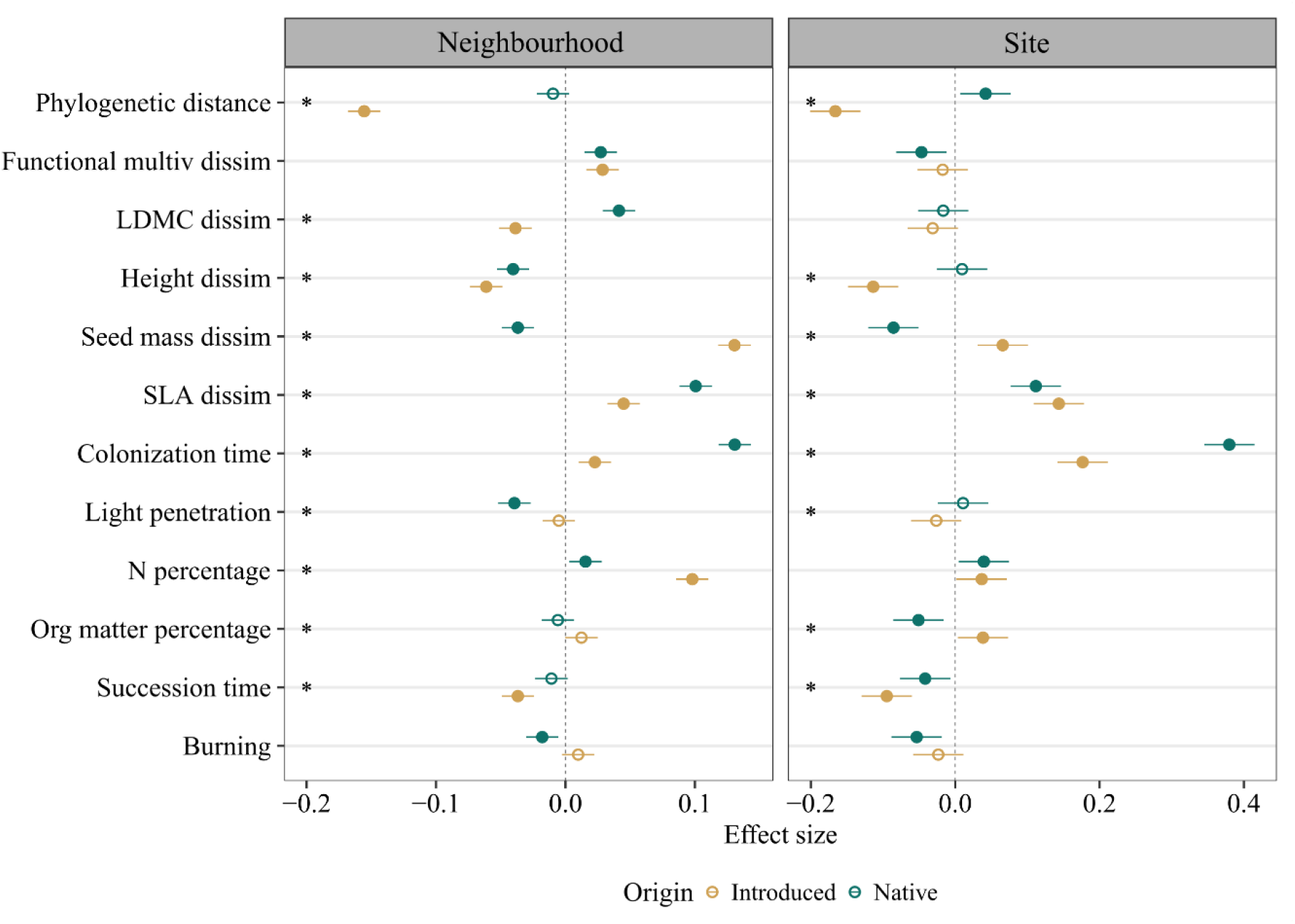
Effect size with 95% confidence intervals of each single explanatory variable included in the final selected model both for neighbourhood (0.5 m^2^) and site spatial scales (~40 m^2^). See Table S2 for further details on model structure and variable transformation. Dark green dots represent native species and ochre dots represent introduced species. Filled dots and empty dots represent statistically significant (p < 0.05) and non-significant effects (p > 0.05), respectively. Asterisk in the left of each panel indicates the presence of statistically significant differences between effect sizes of native and introduced species.

**Figure 5.**
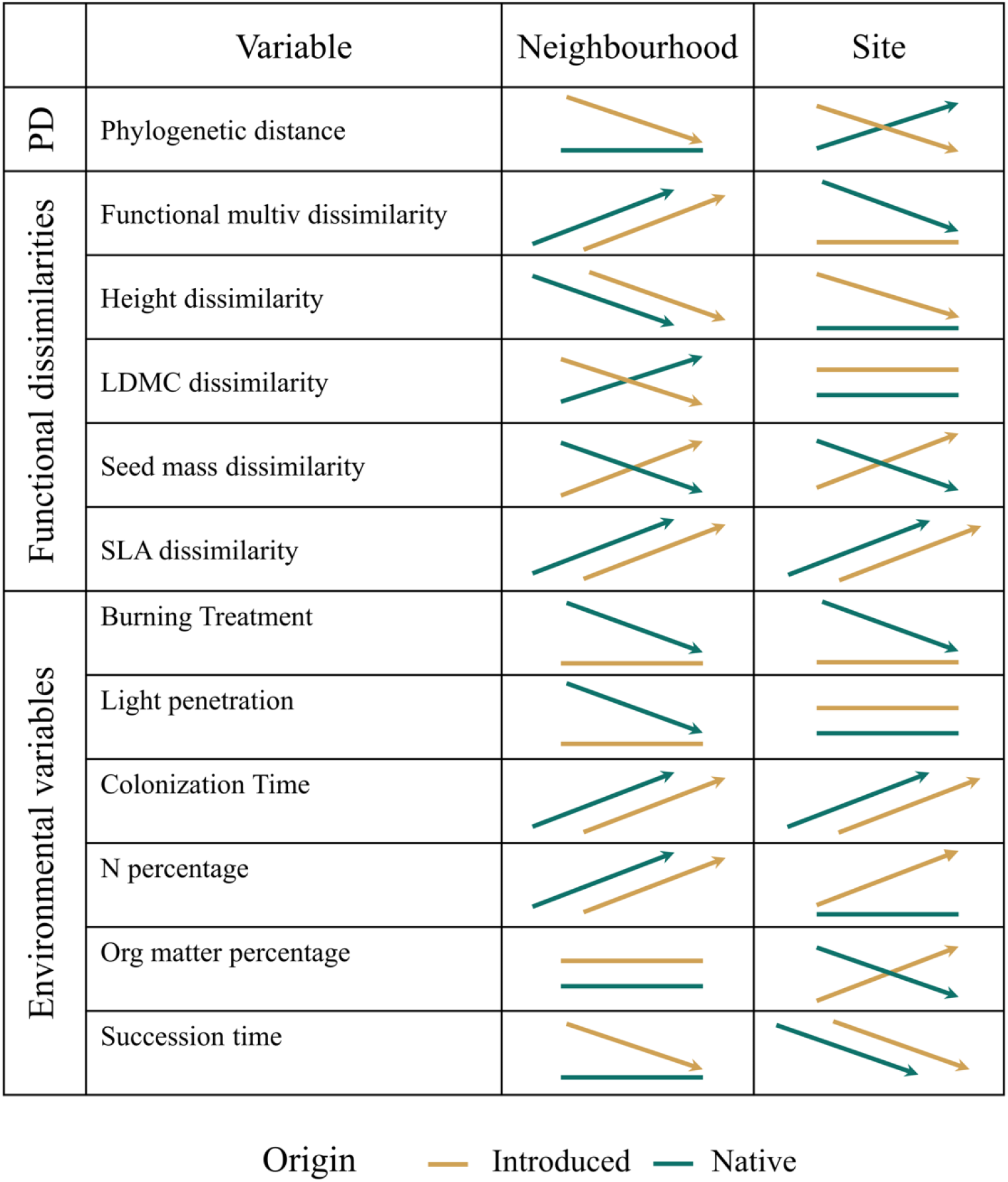
Visual summary of overall relationships between species abundances and the explanatory variables at neighbourhood (5 m^2^) and site (~40 m^2^) scales. Figure is a visual aid for summary purposes only; actual relationships shown in Figs 2-4. For each panel, the X-axis represents an increase in the listed variable, and the Y-axis shows the response of species’ percent cover. The slope of the arrows indicate the direction of observed trends (either positive or negative) but not its magnitude. Flat lines with no arrow heads indicate relationships that were not statistically significant. Dark green colour represents native species and ochre colour introduced species.

Relationships between species abundance and environmental variables were often not statistically significant, particularly at neighbourhood scale (Figs. 4-5). When they were significant, introduced and native species’ relationships usually differed in strength but not in sign (Fig. 4). For example, abundance of both native and introduced species increased with higher soil N concentrations at neighbourhood scales, though introduced species abundance increased more (Fig. S8g). Species abundances increased with colonization time and decreased with succession time both for native and introduced, but introduced species had lower effect sizes than natives in both cases (Fig. 3e-h). In contrast, introduced and native species had opposite relationships with soil organic matter concentration, with introduced species increasing and native species decreasing at the site scale (Figs 4, 5 & S8j, Table S8).

## Discussion

We found that support for the two hypotheses underlying Darwin’s naturalization conundrum varied depending on study spatial resolution and metric used to characterize species differences. Trends associated with multivariate functional dissimilarities were consistent with the limiting similarity hypothesis at the neighbourhood scale (0.5 m^2^) and with the pre-adaptation hypothesis at the site scale (~40 m^2^). Relationships between introduced species abundance and their phylogenetic distance were consistent with the pre-adaptation hypothesis at both scales. Introduced and native species shared similar relationships with multivariate functional dissimilarity, but the relationships between species abundances and phylogenetic distance, ranked univariate functional dissimilarities and environmental drivers differed depending on species biogeographic origin. These last results suggest that, overall, the introduced and native species possessed different life history strategies and competitive abilities.

### 1. Support for limiting similarity and pre-adaptation hypotheses across spatial scales

The different relationships between species abundance and multivariate functional dissimilarity and phylogenetic distance highlight an apparent discrepancy between these two dissimilarity metrics, particularly at finer spatial scales, with functional dissimilarity relationships supporting the limiting similarity hypothesis and phylogenetic distance relationships supporting the pre-adaptation hypothesis (Figs 1,5). Our findings confirm the need to account for both functional and phylogenetic dissimilarities when examining Darwin’s naturalization conundrum (Cadotte et al., 2013; Galland et al., 2019; Gallien & Carboni, 2017; Pinto-Ledezma et al., 2020).

Support for the limiting similarity hypothesis at finer spatial scales, indicated by the positive relationship between abundance and functional dissimilarity, was consistent with the general expectation of greater detection of plant competition at finer spatial scales (Kraft et al., 2015). Similarly, support for the pre-adaptation hypothesis at site scale agreed with the increasing relevance of environmental filtering in community assembly at larger spatial resolutions (Emery & Kellogg, 2007; Ma et al., 2016; Park et al., 2020). While this trend was shared by native and introduced species at the neighbourhood scale, support for the pre-adaptation hypothesis at the site scale was not statistically significant for introduced species in terms of functional dissimilarity. The role of multiple mechanisms in determining invasion may have prevented higher support for pre-adaptation at the site scale (Catford et al., 2019). The fact that the relationship between species abundance and functional dissimilarity differed across spatial scales as small as 0.5 m^2^ and 40m^2^ suggests that spatial scales of tens of metres, frequently used to represent neighbourhood-local conditions (Götzenberger et al., 2012), might be too large to evince limiting similarity. Variation in study spatial scales likely contribute to the mixed conclusions about Darwin’s naturalization conundrum (Cadotte et al., 2018; Catford et al., 2022).

Relationships between species abundance and phylogenetic distance provided support for the pre-adaptation hypothesis at both spatial scales, such that introduced species closely related to co-occurring species had higher abundances (Figs. 2-3). Multiple studies have recently analysed and reviewed the prevalence of limiting similarity and pre-adaptation hypotheses across spatial scales using phylogenetic distances (Cadotte et al., 2018; Carboni et al., 2013; Ma et al., 2016; Park et al., 2020), obtaining higher support for the pre-adaptation hypothesis mainly at larger scales or showing inconclusive results. Given the relatively fine resolution of spatial scales analysed in our study, our results disagree with some of these previous studies by supporting pre-adaptation hypothesis even at fine spatial scales.

Somewhat unexpectedly, we found simultaneous support for the pre-adaptation and limiting similarity hypothesis at the same (neighbourhood) scale when using phylogenetic distance and functional dissimilarity respectively. This pattern might emerge due to: 1) the lack of correlation between phylogenetic and functional measures, due to use of traits with weak phylogenetic signal (Flynn et al., 2011); and 2) that species colonization occurs in a stepwise process where the introduced species more closely-related to natives are able to survive in the environmental conditions of the new range, and then those more functionally similar species from these taxa are removed at finer resolutions due to competitive exclusion (Divíšek et al., 2018). Nonetheless, these interpretations should be considered with caution given the lower effect size of multivariate functional dissimilarity as a predictor of vegetation cover patterns compared to the effect size of phylogenetic distance.

### 2. Differences between native and introduced species

Our study shows that introduced and native species can have similar relationships with some drivers, but distinct or opposite relationships with others. Species found at the edge of multivariate functional trait space reached higher abundance at neighbourhood scale, regardless of whether they were native or introduced, suggesting that limiting similarity is a common driver of community assembly at finer resolutions, regardless of species origin. This result aligns with assertions that native and introduced species have to follow the same set of “rules” to become abundant (Lemoine et al., 2015, 2016). However, introduced and native species did have distinct relationships with many other variables, including phylogenetic distance at the site scale where introduced species were more abundant when they were more closely related to co-occurring species whereas the opposite was true for natives. Our results also revealed that native and introduced species often differed in their functional strategies (e.g., introduced species were more abundant when having heavier seeds while native showed the opposite strategy) and their relationships with environmental conditions (e.g., introduced species were more abundant than natives at higher N soil concentration). These different trends suggest that, depending on biogeographic origin, species can show different responses to ecological forces and possess distinct life history strategies (Zheng et al., 2015), highlighting the value of considering species’ origin when assessing performance and ecological impacts (Buckley & Catford, 2016).

### 3. Ranked differences of univariate traits

Our study revealed that introduced species were more abundant when showing higher ranked values for traits related with leaf acquisitive strategies and seed mass but lower height. Specifically, introduced species’ abundance was related to higher SLA and lower LDMC values relative to the recipient community, suggesting higher rates of resources uptake (Díaz et al., 2016; Reich, 2014; Wright et al., 2004) and supporting common findings where invaders with faster leaf economics are more successful (Catford et al., 2019; Van Kleunen et al., 2010). Although the correlation between introduced species abundance and seed mass varies (Catford et al., 2016; Pyšek & Richardson, 2008), including at Cedar Creek (Catford & Jones, 2019; Pinto-Ledezma et al., 2020), we found that introduced species with heavier seeds were more abundant at both spatial scales (though native species showed the opposite trend). Species with heavier seeds tend to have higher establishment and survival in resource-limited environments than species with light seeds (Tilman, 1994), which implies that the introduced species in our study not only had fast resource-acquisitive strategies but also had seeds more resistant to low resource conditions. This combination of strategies could give them an advantage over co-occurring native species.

Introduced species at both scales, and natives at the neighbourhood scale, were more abundant when they were shorter than other plant species within the community. This finding contradicts the usual expectation that taller plants will be more abundant than shorter plants because of their superior competitive ability for light (Pyšek & Richardson, 2008). However, previous studies carried out at Cedar Creek found no conclusive differences in height ranked differences between native and introduced species (Pinto-Ledezma et al., 2020), confirming the low relevance of light competition in the study grasslands (Tilman, 1990).

Contrary to multivariate functional dissimilarities, where introduced and native species had similar trends, the ranked univariate dissimilarities revealed different patterns depending on species origin. Ranked indexes emphasize the hierarchical position of each species on each trait gradient, and thus can better reflect competitive capacities. In contrast, multivariate dissimilarity shows the absolute distance between a given species’ trait combination and the community trait centroid, independently of the species exact ranked position in the 3D functional space, thus better representing use of different niches (Catford et al., 2019; Gallien & Carboni, 2017). Our results, therefore, suggest that although all species benefit from using different niches and avoiding overlap, introduced and native species tend to occur at different positions along trait gradients and possess different life history strategies.

### 4. Land-use history and environmental variables modulate introduced species abundance

Species invasion is affected by land use history and environmental variables (González-Moreno et al., 2014; Theoharides & Dukes, 2007). Our study indicated the introduced species had greater capacity to compete for nutrients in resource limited environments (Matzek, 2011), as introduced species showed higher abundances than native along soil nitrogen and organic matter concentrations. Time since species colonization also had a positive effect on species abundance, though introduced species increased their abundance faster and native species reached higher abundances with time, implying different strategies in the competition-colonization trade-off (Calcagno et al., 2006). Conversely, introduced and native species reduced their abundances as time since abandonment increased (at both spatial scales), which is consistent with a reduction in resource availability and increase in competition with time since abandonment (Catford et al., 2012; Kneitel & Perrault, 2006). A decline in abundance with increasing succession time was particularly pronounced for introduced species, which were more abundant than natives in early succession and less abundant than natives in late succession. Similar trends have been found in old and young forests (Lemoine et al., 2016).

## 5. Conclusion

By examining hypotheses underpinning Darwin’s naturalization conundrum across two spatial scales in a long-term field experiment, we found evidence for the pre-adaptation hypothesis (environmental filtering) at the site scale (~40 m^2^). This suggests that environmental heterogeneity strongly influences grassland community assembly even at very small spatial scales in grasslands. Importantly, this sort of scale (40 m^2^) is frequently used as the smaller “local” scale in studies examining the conundrum, based on the premise that it can reveal effects of competition and limiting similarity. Thus, many of the inconsistences around support for the limiting similarity hypothesis could arguably stem from inappropriate selection of study spatial scales. Our results also revealed how support for hypotheses can depend on the variables used to examine them.

Multivariate functional dissimilarity and phylogenetic distance each supported different hypotheses at neighbourhood scale, even though these two variables are often considered to be proxies for one another. Thus, we highlight that i) selecting inaccurate spatial scales that fail to capture the corresponding the target community assembly process (competition or environmental filtering), and ii) the independent use of either functional or phylogenetic approaches might cause the apparent context dependence and lack of consensus surrounding Darwin’s naturalization conundrum. We also found that introduced and native species frequently had different relationships with environmental covariates. This suggests that species’ responses to environmental conditions and functional strategies for reaching high abundance can vary depending on species origin.

## Acknowledgements

We are very grateful to David Tilman as responsible of E-014 experiment at Cedar Creek as well as to all students, interns and researchers for taking care of and surveying the experiment over several decades. The research underlying this study has been supported by the European Research Council (ERC) under the European Union’s Horizon 2020 research and innovation programme (grant agreement No. [101002987]). The authors also acknowledge financial support from the University of Graz. Data collection was supported by the NSF LTER program, including DEB-8114302, DEB-8811884, DEB-9411972, DEB-0080382, DEB-0620652, and DEB-1234162, and by Cedar Creek Ecosystem Science Reserve and the University of Minnesota.

## Author contributions

Maria A. Perez-Navarro conceptualized the original idea, led the data curation, did the formal analyses, and wrote the first draft. Adam Clark advised for data curation and analytical models. Josh Brian and Harry Shepherd advised on figure representation and analytical models. Jane Catford led funding acquisition, conceptualized the idea, and advised on figure representation and analytical models. All authors contributed to write subsequent versions of the manuscript.

## Data Availability Statement

Most dataset used in this paper are already publicly available. Field data for experiment E-014 of Cedar Creek Ecosystem Science Reserve can be downloaded at https://www.cedarcreek.umn.edu/research/data/methods?e014 under request. Phylogenetic information is already included in V.phylomaker package available at https://github.com/jinyizju/V.PhyloMaker. Finally, though trait data used in this manuscript are already included in previous studies (Cadotte et al., 2009; Catford et al., 2019; Willis et al., 2010 and experiment E-133), the authors agree to make publicly available the final trait database used after filtering and homogenising these databases.

