## Supporting Information for "Influence of functional and phylogenetic dissimilarity on exotic plant invasion depends on spatial scale"

### Spatial scale determines whether functional and phylogenetic dissimilarity helps or hinders exotic plant invasion

#### Supporting Information

#### Appendix S1. Further details on field sampling and data preparation

**Estimation of soil N, C, organic matter percentage and light penetration**

Soil organic matter, nitrogen and carbon content were measured at plot level at the beginning of the surveys in 1983, and soil nitrogen and carbon were measured every 6 years until 2011. Both nitrogen, carbon and organic matter concentrations were obtained as dry mass percentage, from one soil core of 25 mm x 100 mm located at the centre of each plot. Chemical analyses were conducted using a Carlo Urba NA 1500 elemental analyser (Knops & Tilman 2000; Knops & Bradley 2009). Light penetration was recorded in 1984, 1989 and 2016 at transect and plot level, respectively. In each case a pair of light meter readings were taken, one at ground level and one above the vegetation. Then, the light penetration percentage was obtained as light at ground level/light above vegetation*100. Since some plots and transects lacked light and soil data, we interpolated plot and transect level data when required to fill the gaps. When soil and light data were missing at plot level, average data from the rest of plots of the same transect were used. When transect data were missing, average data from the rest of transects within the same field were used. Since soil and light sampled years do not match the exact years of plant abundance surveys, we have averaged temporal soil and light data at plot or transect level.

**Estimation of Minimum Colonisation Time**

Minimum colonisation time in the community was estimated as the as the difference between the year that a species was observed in a given plot and the year when the species first colonised the plot. The year of first colonisation was estimated as the midpoint year between the year the species was observed the first time in the plot and the previously sampled year in the plot, when the species was still not present. For example, if a given species appeared in a plot for the first time in 1989 and the plot was previously sampled in 1983, we would infer 1986 as colonisation year. We used this approach instead of using the first-observed record as colonisation year (Castro et al., 2005; Wilson et al., 2007) since not all plots were sampled with the same temporal recurrence, so the first-observed year could be less precise in plot sampled with less frequency. However, when the species was first observed in the plot the first year the plot was sampled, it was not possible to determine the species colonisation year basing on the previous sampled year. In this case, we followed one of the next two approaches to estimate colonisation year: 1) if the plot was abandoned shortly before the beginning of the surveys (i.e. less than 6 years before, which is the average gap between surveys), the colonisation year was estimated as the midpoint between year of species appearance in the plot and year of field abandonment; and 2) if the plot was abandoned more than 6 years before the beginning of the surveys, we estimated the colonisation year as the year of plot abandonment plus the species' colonisation delay. The colonisation delay is defined as the period of time that the species delay appearing after crop abandonment. We calculated it by estimating the difference between the species first-colonisation and the year of abandonment in those cases where it was possible to infer the colonisation year with high precision (i.e., the species appeared after the first survey, or the field was abandoned short before the appearance of the species in the plot). Then, used the mode (the most likely value) of this distribution of delay values as the species representative delay in colonisation. We used the mode instead the average or median since median or average values may not match a likely colonisation value in case of bi or multimodal distributions (Appendix S1-Fig. 1).

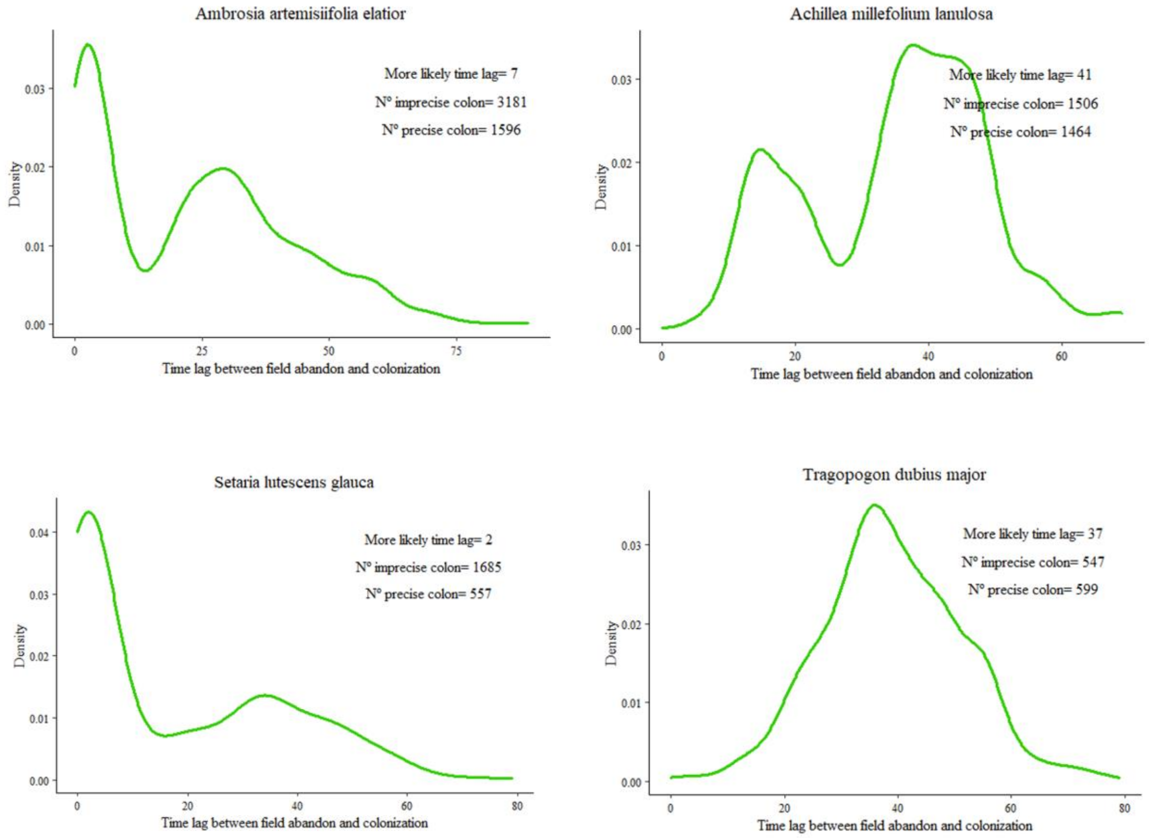

**Appendix S1-Figure 1.** Density plot of time lapse between cropland abandon and first documented occurrence in a given plot for different species. "Imprecise colon" referrer to the number of plots which appeared on the first sampling year and where the time lag between colonisation and abandon is higher than the usual lag between sampling years. "Precise colon" referrer to the number of plots which appeared after the first sampling year or appeared on the first sampled year but the time lag between colonisation and abandon is higher equal or lower the usual lag between sampling years. "More likely time lag" refers to the most probable number of years that a given species need to colonize. Note that the ratio between the number of precise and imprecise colonisation cases suggests the robustness of imputation of time lag after abandon. The higher number of imprecise colonisation cases/number precise colonisation cases the less plausibility and robustness of the imputed value.

#### Appendix S2. Further details on trait data

Functional trait data for the study species were compiled from trait databases of five different studies carried out at Cedar Creek Ecosystem Science Reserve (Cadotte et al., 2009; Catford et al., 2019; Willis et al., 2010 and experiment e133). We compiled a total of 56 quantitative functional traits across species and datasets (Fig. S2 and S3). For each study dataset we averaged trait data at species level when more than one individual per species was sampled. Among the 56 total functional traits we selected seven that had the highest coverage across species: specific leaf area (SLA), plant height, leaf area, leaf dry matter content (LDMC), leaf fresh mass, leaf dry mass (all of them with data for more than 60% of the species) and seed mass (with data for 55% of the species). Although there were some missing values in the trait-species matrix, native and introduced species were equally well represented (Fig. S2). For taxa that could only be identified to genus level, we used average trait data across the species of the corresponding genus present in the study site. When a species had data for a given functional trait from more than one dataset, we choose the trait dataset following the next quality criteria: Catford et al. 2019>Cadotte et al. 2009>Willis et al. 2010> e133, attending to the number of replicates used to obtain the functional trait value (Catford between 3 and 15 replicates, Cadotte et al. 2009 10 replicates, Willis et al. 2010, and e133 3 replicates), the presence of interpolated data from other datasets (e133), and the publication status (published vs unpublished in peer-review journals). We did not average data across datasets to ensure the prevalence of maximum quality datasets. Finally, most of the trait data came from Catford et al. 2019 dataset.

#### Appendix S3. Appendix References

Cadotte, M.W., Cavender-Bares, J., Tilman, D. & Oakley, T.H. (2009). Using phylogenetic, functional and trait diversity to understand patterns of plant community productivity. *PLoS One*, 4, 1–9.

Catford, J.A., Smith, A.L., Wragg, P.D., Clark, A.T., Kosmala, M., Cavender-Bares, J., *et al.* (2019). Traits linked with species invasiveness and community invasibility vary with time, stage and indicator of invasion in a long-term grassland experiment. *Ecol. Lett.*, 22, 593–604.

Knops, J.M.H. & Bradley, K. (2009). Soil Carbon and Nitrogen Accumulation and Vertical Distribution across a 74-Year Chronosequence. *Soil Sci. Soc. Am. J.*, 73, 2096–2104.

Knops, J.M.H. & Tilman, D. (2000). Dynamics of soil nitrogen and carbon accumulation for 61 years after agricultural abandonment. *Ecology*, 81, 88–98.

Willis, C.G., Halina, M., Lehman, C., Reich, P.B., Keen, A., McCarthy, S., *et al.* (2010). Phylogenetic community structure in Minnesota oak savanna is influenced by spatial extent and environmental variation. *Ecography (Cop.).*, 33, 565–577.

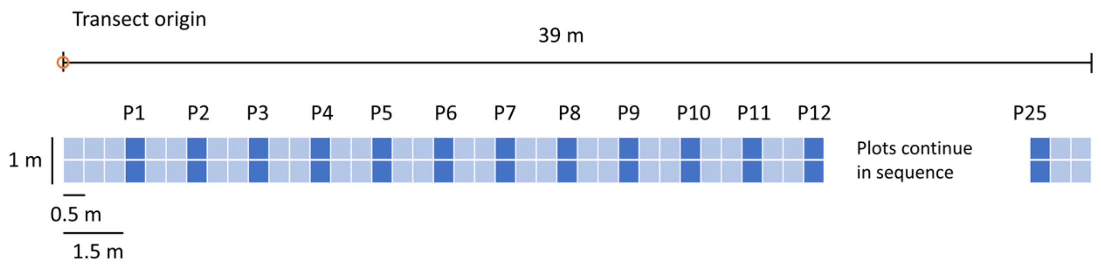

**Figure S1.** Transect spatial structure. P1 to P25 represent each of the plots nested within the transect. Dark blue represents the sampled plots and clear blue colour the non-sampled surface between plots. Figure adapted from Cedar Creek web page:

(https://www.cedarcreek.umn.edu/research/data/methods?e014)

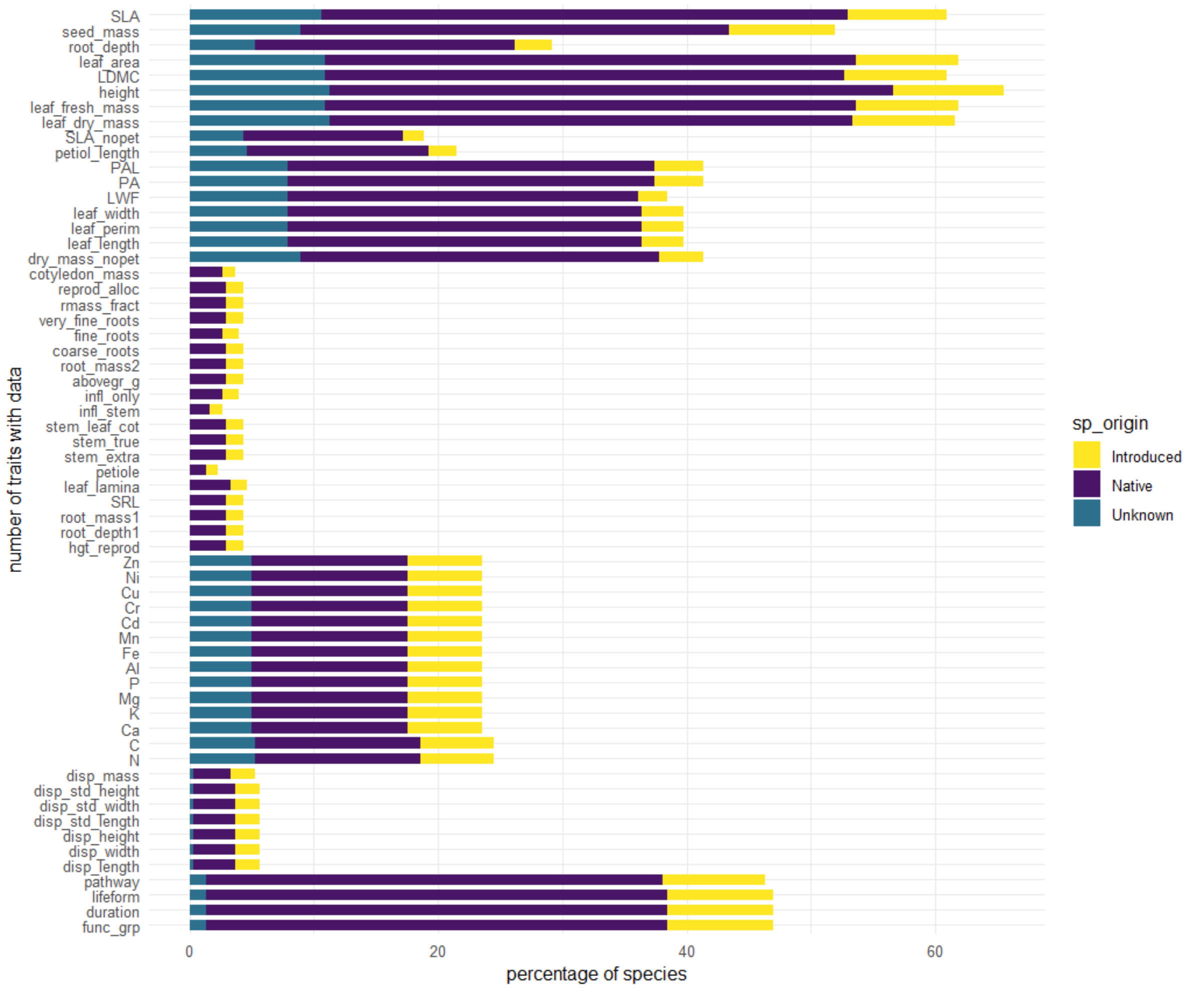

**Figure S2.** Percentage species with data for each trait variable per origin status. Different colours represent the species origin (yellow – introduced species, purple – native species and blue – species with unknown origin). Total number of species is 303.

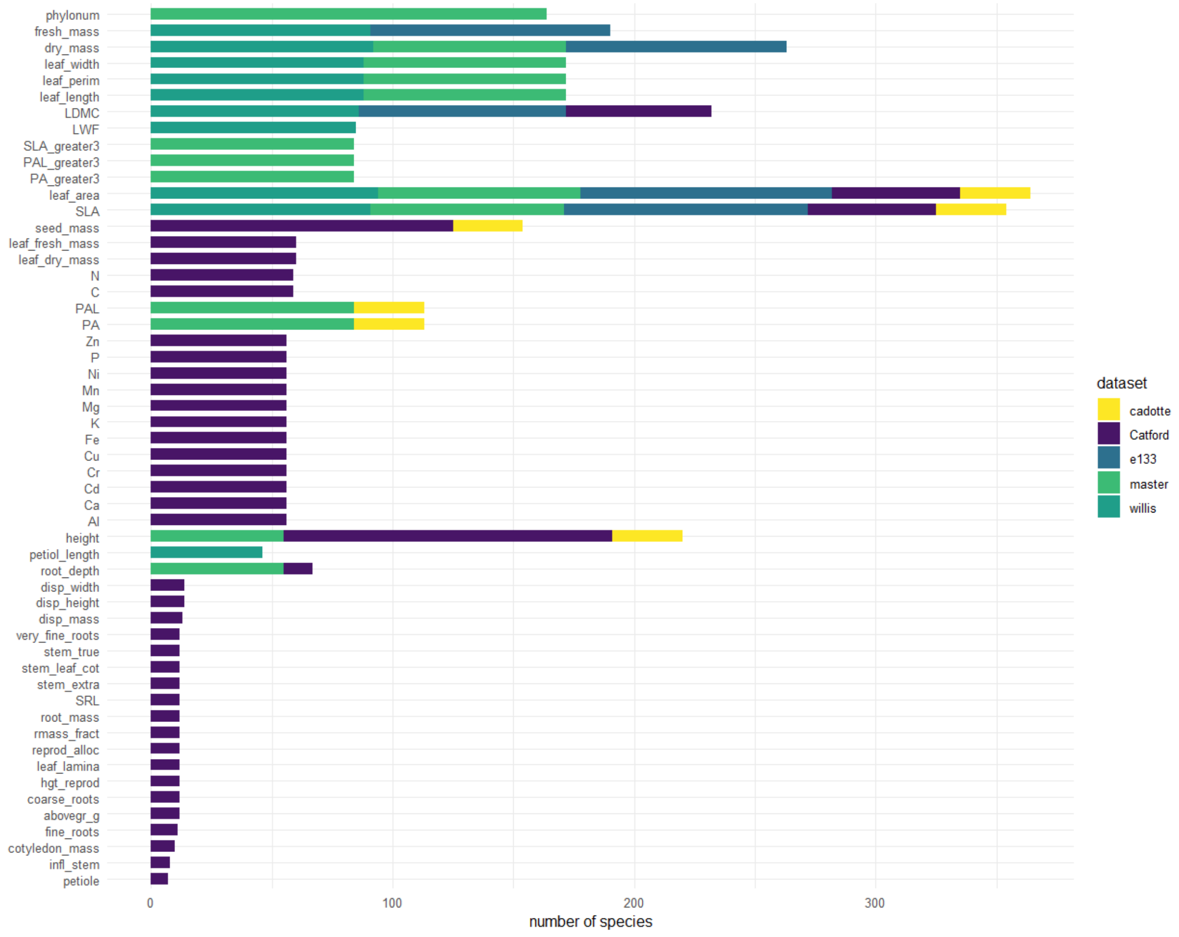

**Figure S3.** Number of species with trait data regarding to source trait dataset. In yellow Cadotte et al. 2009, in purple Catford et al. 2019, in dark blue e133 experiment data, in green unpublished data from a master thesis supervised by Willis, and in blue-green Willis et al. 2010. Please consider that a same species could have data from more than one dataset, so the sum of all the covered species by different datasets may be higher than the total species richness of the study (303 species).

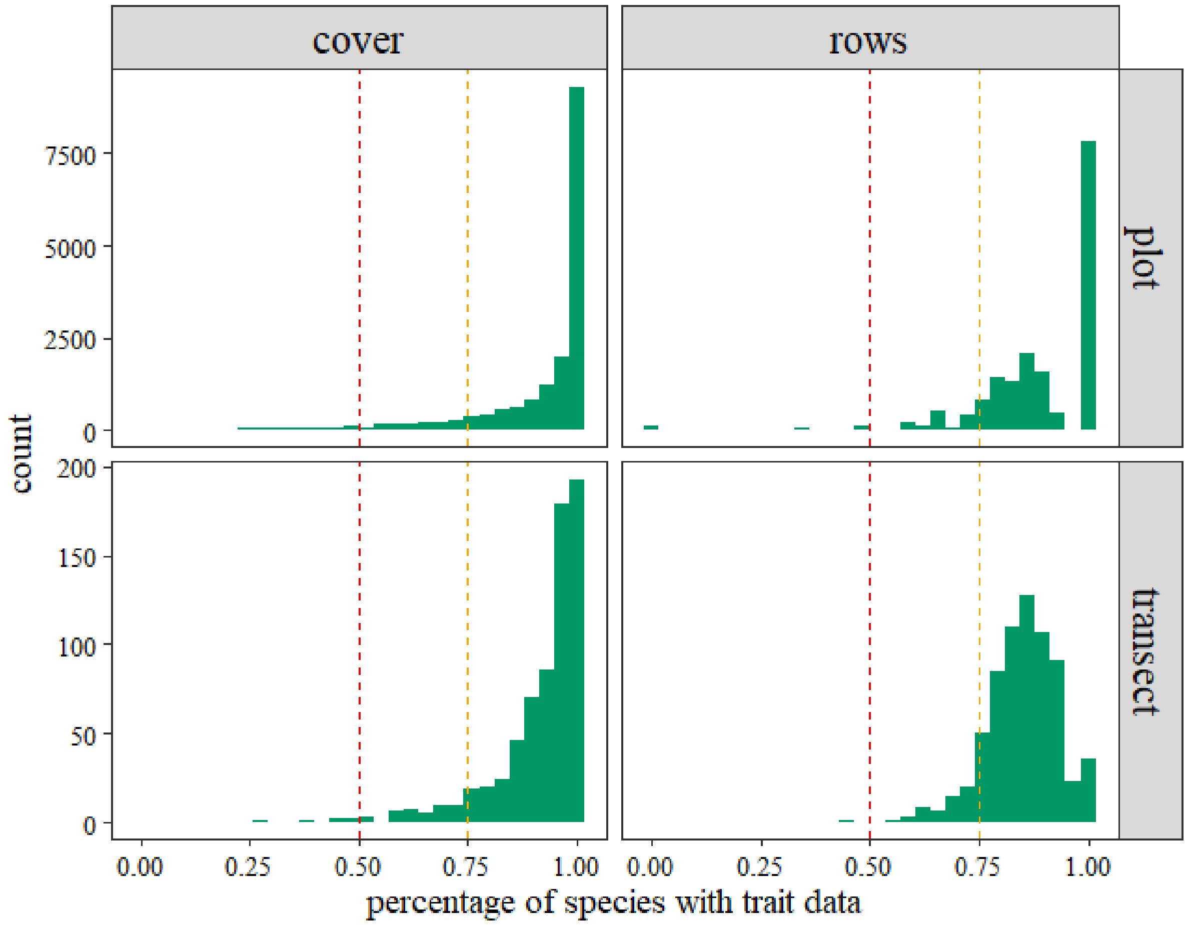

**Figure S4.** Proportion of species with functional trait data at plot and transect level from the total number of species present within each plot or transect. Left column shows frequencies regarding plant cover (i.e., Proportion of plot/transect cover with trait data) and right column shows frequencies regarding number of species (i.e., Table rows, at species and transect level). Red dashed line shows the threshold of 50% of species/cover having functional data per sampling unit, and orange dashed line shows the threshold of 75% of species/cover with functional data per sampling unit.

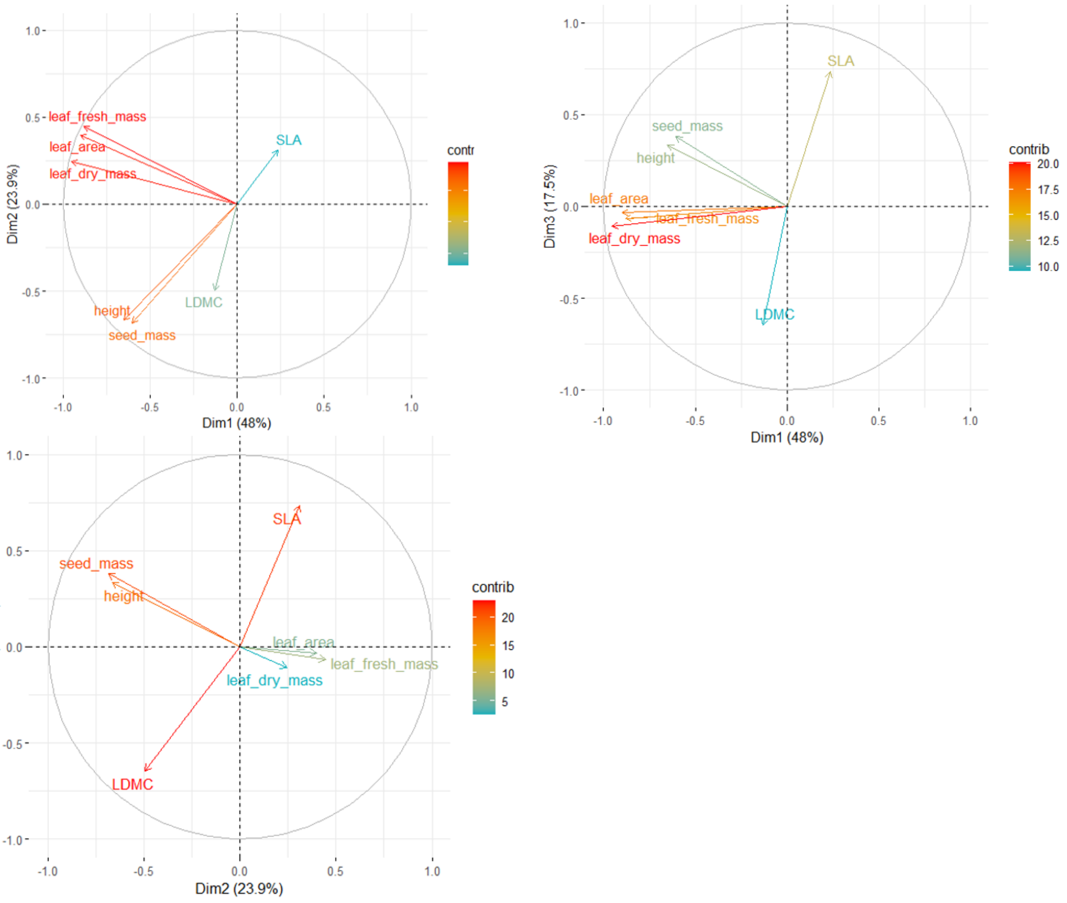

**Figure S5.** Correlation circles obtained from PCA for the 7 selected functional traits: SLA, seed mass, leaf area, LDMC, height, leaf fresh mass, and leaf dry mass. The three first axis accumulated the 75.37% of total variability. Top left panel represents the orthogonal distribution of functional traits in the bidimensional space of PCA dimensions 1 and 2; top right panel represents the orthogonal distribution of functional traits in the bidimensional space of PCA dimensions 1 and 3; and bottom left panel represents the orthogonal distribution of functional traits in the bidimensional space of PCA dimensions 2 and 3. Variables' colour represents the percentage of each variable contribution to the PCA.

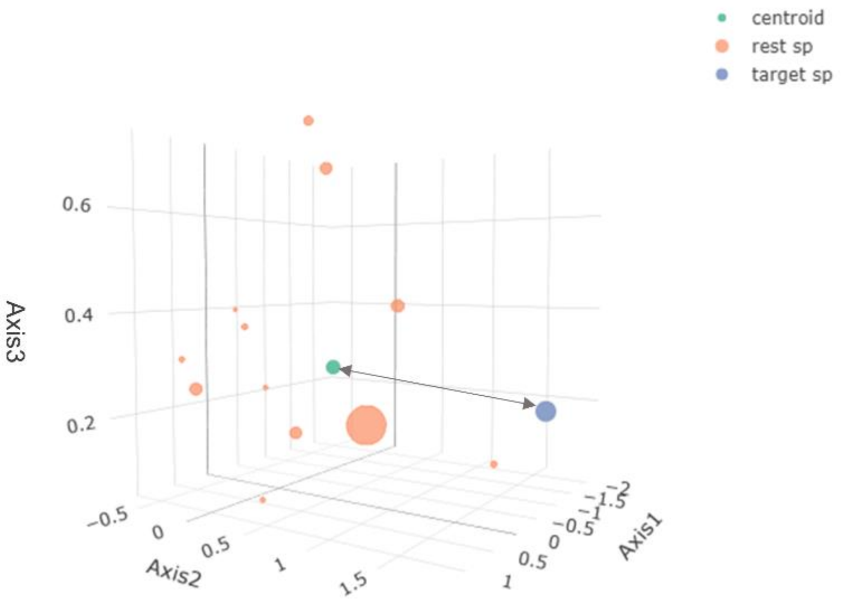

**Figure S6.** Diagram representing the approach used to measure the functional dissimilarity in the three-dimensional (3D) functional multivariate trait space built with the seven selected functional variables: SLA, seed mass, leaf area, LDMC, height, leaf fresh mass, and leaf dry mass. Axis 1 explains 45.4% of the total traits' variance, while Axis 2 and Axis 3 explain 19.8% and 14.4% of the total variance, respectively. This diagram is an example built with species data from transect A, field 39 of 2016 sampled year. Blue point represents the target species position (Andropogon gerardii) in the functional multivariate space, while orange points represent the position of the rest of species of the same community. The green dot represents the 3D position of the community centroid of the transect community (estimated as the centre of mass of the rest of the species but the target species). Point size represents the species relative abundance within the community. Multivariate functional dissimilarity was estimated following the "centroid distance method", in which we measured the distance between the target species functional position in 3D and the position of the community centroid.

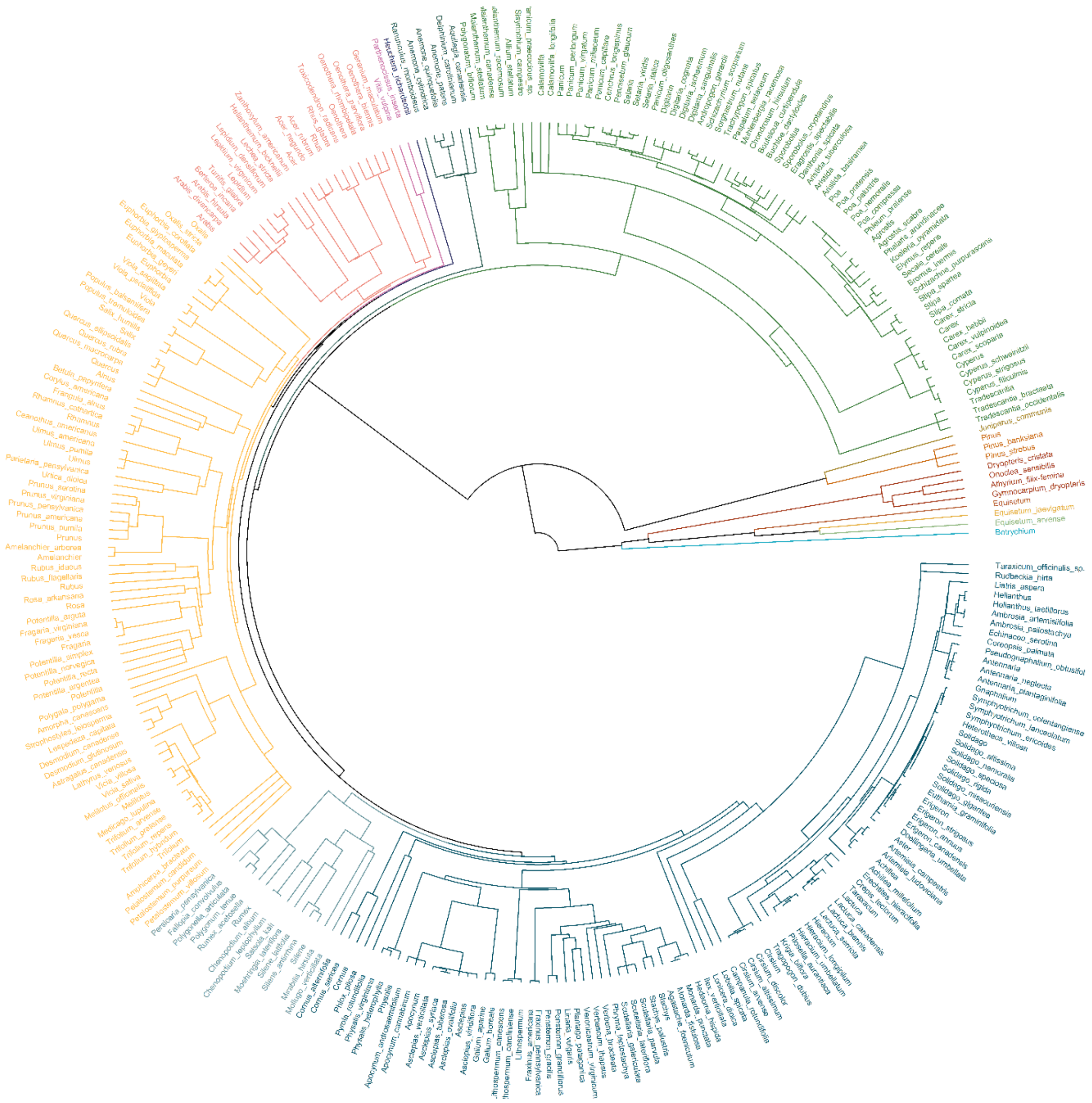

**Figure S7.** Phylogenetic tree for all the 303 species included in e014 Cedar Creek experiment database based on Smith and Brown 2018 and Zanne et al. (2014) phylogeny. Given the limited size of word document, please visit separated supplemenatry phylogenetic circle in pdf format for consulting specific species names.

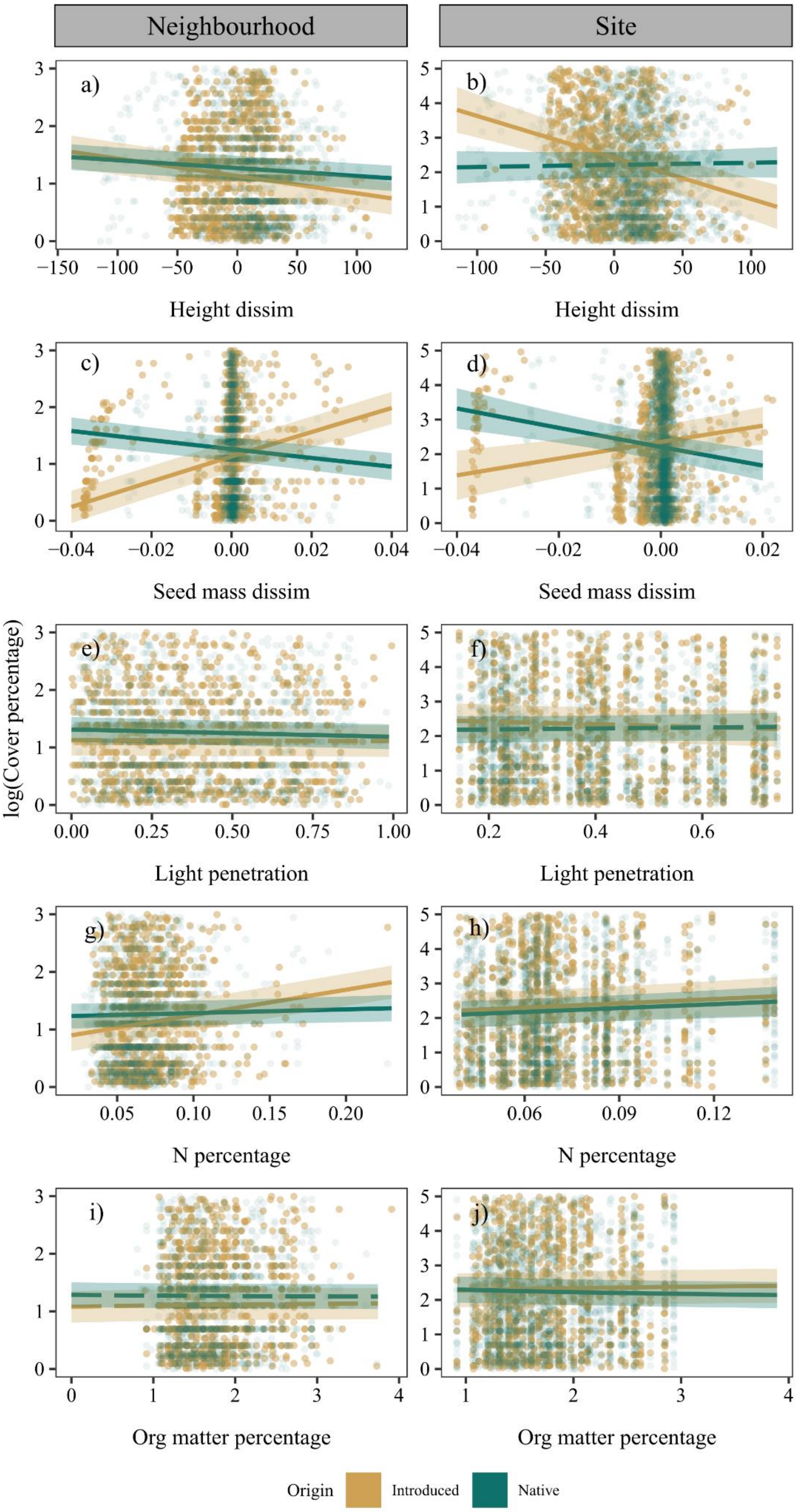

Neighbourhood
Site
a)
b)
c)
d)
e)
f)
g)
h)
i)
j)
Height dissim
Height dissim
Seed mass dissim
Seed mass dissim
Light penetration
Light penetration
N percentage
N percentage
Org matter percentage
Org matter percentage
log(Cover percentage)
Origin
Introduced
Native

**Figure S8.** Regression plots corresponding to final selected model for both neighbourhood and site spatial scales. Y axes represent log of species cover percent. X axes correspond to each of the explanatory variables. Bluegreen color represents native species populations and brown color represents native species populations. Solid and dashed lines represent significant ($p < 0.05$) and not significant effects ($p > 0.05$), respectively. Plots only includes 5% of species x plot data used in models at neighbourhood level and 25% of species x transect data at site level, in order to reduce the number of points (to 5000 each) and improve visualization.

**Table S1.** Definitions and units of measurement of traits included in the study.

| Trait | Unit | Comments |
|---|---|---|
| fresh_mass | g | Fresh lamina mass including petiole (g) |
| dry_mass | g | Dry lamina mass including petiole (g) |
| LDMC | mg/g | Lamina dry matter content including petiole (mg/g) |
| Leaf area | $mm^2$ | fresh leaf area (scanned) |
| SLA | $mm^2/mg$ | area/(dry_mass*1000) |
| plant height | cm | vegetative height |
| seed mass | g | seed mass for a seed sample/number of seeds |

**Table S2.** Variables included in each of the 6 tested alternative models. Variables were selected after discarding regression multicollinearity. Thus, carbon percentage, dissimilarity in leaf area, dissimilarity in dry matter content and dissimilarity in leaf fresh mass are not included in the table as they were discarded from models including environmental variables and univariate functional dissimilarity, respectively, due to high collinearity. Green colour shading represents environmental variables, orange colour shading represent functional trait variables and yellow colour represents phylogenetic variables. Final selected model corresponds to model 4.

| Variable code | Variable | Transformation | Model 1 | Model 2 | Model 3 | Model 4 | Model 5 |
|---|---|---|---|---|---|---|---|
| PD | Phylogenetic distance | scaled | x | x | x | x | x |
| MFD | Multivariate functional distance | log | x | x | x | x | x |
| UFD_sm | Seed mass dissimilarity | log | | | x | x | x |
| UFD_sla | specific leaf area dissimilarity | scaled | | | x | x | x |
| UDF_ldmc | leaf dry matter content dissimilarity | scaled | | | x | x | x |
| UFD_h | plant height dissimilarity | scaled | | | x | x | x |
| ENV_tab | Time since abandonment | scaled | | x | | x | x |
| ENV_mrt | Minimum Colonisation Time | log | | x | | x | x |
| ENV_n | Soil nitrogen content | | | x | | x | x |
| ENV_c | Soil carbon content | | | | | | x |
| ENV_om | Soil organic matter content | log | | x | | x | x |
| ENV_b | Burning treatment | | | x | | x | x |
| ENV_lp | Light penetration | | | x | | x | x |
| ENV_tab_int | Time since abandonment Interacting with each of the rest explanatory variables | scaled | | | | | x |

**Table S3.** Model summary for each of the fitted models at neighbourhood level (plot scale). Table shows: $R^2$ marginal ($R^2$ m), $R^2$ conditional ($R^2$ c), Akaike Information Criterion (AIC) and the difference between each model AIC, the difference between model AIC and the minimum AIC value across models (AIC dif), Bayesian Information Criterion (BIC) and BIC difference between each model BIC and the lowest BIC found across models. Final selected model and further results corresponds to model 4.

| **MODEL** | **$R^2$M** | **$R^2$C** | **$AIC_C$** | **$AIC_C$_DIF** | **BIC** | **BIC_DIF** |
|---|---|---|---|---|---|---|
| **MOD 1** | 0.005 | 0.378 | 273691.58 | 25847.84 | 273806.44 | 25659.51 |
| **MOD 2** | 0.027 | 0.313 | 248893.67 | 1049.94 | 249121.48 | 974.55 |
| **MOD 3** | 0.049 | 0.413 | 272665.56 | 24821.83 | 272857.00 | 24710.07 |
| **MOD 4** | 0.081 | 0.358 | 247843.73 | **0.00** | 248147.47 | **0** |
| **MOD 5** | 0.05 | 0.355 | 248164.90 | 321.17 | 248468.60 | 321.40 |

**Table S4.** Model summary for each of the fitted models at site level (transect scale). Table shows: $R^2$ marginal ($R^2$ m), $R^2$ conditional ($R^2$ c), Akaike Information Criterion (AIC) and the difference between each model AIC, the difference between model AIC and the minimum AIC value across models (AIC dif), Bayesian Information Criterion (BIC) and BIC difference between each model BIC and the lowest BIC found across models. Final selected model and further results corresponds to model 4.

| MODEL | $R^2$M | $R^2$C | AICC | AICC_DIF | BIC | BIC_DIF |
|---|---|---|---|---|---|---|
| **MOD 1** | 0.015 | 0.395 | 46634.44 | 4432.53 | 46717.11 | 4284.61 |
| **MOD 2** | 0.065 | 0.431 | 42535.68 | 333.77 | 42706.88 | 274.38 |
| **MOD 3** | 0.135 | 0.489 | 46219.40 | 4017.49 | 46369.61 | 3937.11 |
| **MOD 5** | 0.175 | 0.495 | 42201.91 | **0.00** | 42432.50 | **0.00** |
| **MOD 6** | 0.078 | 0.447 | 42557.39 | 230.58 | 42787.97 | 355.47 |

**Table S5.** Model results for the finally selected model at neighbourhood spatial scale including variables slopes of each species origin (native or introduced) separately. Bold pvalue highlits variables's slope significantly different from 0 slope (pv<0.05).

| **Variable** | **origin** | **slope** | **SE** | **df** | **z.ratio** | **p.value** |
|---|---|---|---|---|---|---|
| **Succession time** | Introduced | -0.010 | 0.002 | Inf | -5.757 | **<0.0001** |
| **Succession time** | Native | -0.003 | 0.002 | Inf | -1.699 | 0.089 |
| **Minimum colonisation time** | Introduced | 0.001 | <0.0001 | Inf | 3.555 | **<0.0001** |
| **Minimum colonisation time** | Native | 0.006 | <0.0001 | Inf | 20.464 | **<0.0001** |
| **Phylogenetic distance** | Introduced | -0.002 | <0.0001 | Inf | -24.337 | **<0.0001** |
| **Phylogenetic distance** | Native | <0.0001 | <0.0001 | Inf | -1.509 | 0.131 |
| **Functional multivariate dissimilarity** | Introduced | 0.022 | 0.005 | Inf | 4.489 | **<0.0001** |
| **Functional multivariate dissimilarity** | Native | 0.016 | 0.004 | Inf | 4.262 | **<0.0001** |
| **N percentage** | Introduced | 4.398 | 0.287 | Inf | 15.336 | **<0.0001** |
| **N percentage** | Native | 0.652 | 0.269 | Inf | 2.422 | **0.015** |
| **Org matter percentage** | Introduced | 0.012 | 0.006 | Inf | 1.935 | 0.053 |
| **Org matter percentage** | Native | -0.006 | 0.006 | Inf | -0.921 | 0.357 |
| **Light penetration** | Introduced | -0.018 | 0.023 | Inf | -0.808 | 0.419 |
| **Light penetration** | Native | -0.128 | 0.021 | Inf | -6.175 | **<0.0001** |
| **SLA dissimilarity** | Introduced | 0.006 | 0.001 | Inf | 7.033 | **<0.0001** |
| **SLA dissimilarity** | Native | 0.015 | 0.001 | Inf | 15.748 | **<0.0001** |
| **LDMC dissimilarity** | Introduced | -0.001 | <0.0001 | Inf | -6.047 | **<0.0001** |
| **LDMC dissimilarity** | Native | 0.001 | <0.0001 | Inf | 6.469 | **<0.0001** |
| **Height dissimilarity** | Introduced | -0.003 | <0.0001 | Inf | -9.583 | **<0.0001** |
| **Height dissimilarity** | Native | -0.001 | <0.0001 | Inf | -6.335 | **<0.0001** |
| **Seed mass dissimilarity** | Introduced | 21.783 | 1.066 | Inf | 20.437 | **<0.0001** |
| **Seed mass dissimilarity** | Native | -7.857 | 1.364 | Inf | -5.759 | **<0.0001** |
| **Burning treatment** | Introduced | 0.035 | 0.023 | Inf | 1.536 | 0.125 |

| | | | | | | |
|---|---|---|---|---|---|---|
| **Burning treatment** | Native | -0.063 | 0.022 | Inf | -2.810 | **0.005** |

**Table S6.** Model results for the finally selected model at site spatial scale including variables slopes of each species origin (native or introduced) separately. Bold pvalue highlights variables' slopes significantly different from 0 slope (pv<0.05).

| Variable | origin4 | slope | SE | df | z.ratio | p.value |
|---|---|---|---|---|---|---|
| **Succession time** | Introduced | -0.015 | 0.003 | Inf | -5.326 | **<0.0001** |
| **Succession time** | Native | -0.006 | 0.003 | Inf | -2.340 | **0.019** |
| **Minimum colonisation time** | Introduced | 0.013 | 0.001 | Inf | 9.920 | **<0.0001** |
| **Minimum colonisation time** | Native | 0.018 | 0.001 | Inf | 21.335 | **<0.0001** |
| **Phylogenetic distance** | Introduced | -0.006 | 0.001 | Inf | -9.333 | **<0.0001** |
| **Phylogenetic distance** | Native | 0.001 | <0.0001 | Inf | 2.366 | **0.018** |
| **Functional multivariate dissimilarity** | Introduced | -0.046 | 0.047 | Inf | -0.976 | 0.329 |
| **Functional multivariate dissimilarity** | Native | -0.070 | 0.027 | Inf | -2.624 | **0.009** |
| **N percentage in soil** | Introduced | 4.126 | 1.999 | Inf | 2.064 | **0.039** |
| **N percentage in soil** | Native | 3.765 | 1.689 | Inf | 2.229 | **0.026** |
| **Org matter percentage in soil** | Introduced | 0.045 | 0.021 | Inf | 2.161 | **0.031** |
| **Org matter percentage in soil** | Native | -0.055 | 0.019 | Inf | -2.859 | **0.004** |
| **Light penetration** | Introduced | -0.390 | 0.264 | Inf | -1.475 | 0.140 |
| **Light penetration** | Native | 0.140 | 0.230 | Inf | 0.607 | 0.544 |
| **SLA dissimilarity** | Introduced | 0.043 | 0.005 | Inf | 8.065 | **<0.0001** |
| **SLA dissimilarity** | Native | 0.029 | 0.005 | Inf | 6.276 | **<0.0001** |
| **LDMC dissimilarity** | Introduced | -0.002 | 0.001 | Inf | -1.755 | 0.079 |
| **LDMC dissimilarity** | Native | -0.001 | 0.001 | Inf | -0.934 | 0.351 |
| **Height dissimilarity** | Introduced | -0.012 | 0.002 | Inf | -6.385 | **<0.0001** |
| **Height dissimilarity** | Native | 0.001 | 0.001 | Inf | 0.533 | 0.594 |
| **Seed mass dissimilarity** | Introduced | 23.719 | 6.421 | Inf | 3.694 | **<0.0001** |
| **Seed mass dissimilarity** | Native | -27.297 | 5.681 | Inf | -4.805 | **<0.0001** |

| | | | | | | |
|---|---|---|---|---|---|---|
| **Burning treatment** | Introduced | -0.058 | 0.044 | Inf | -1.317 | 0.188 |
| **Burning treatment** | Native | -0.102 | 0.034 | Inf | -2.997 | **0.003** |

**Table S7.** Differences in slope at neighbourhood spatial scale between introduced and native species slopes for each variable (introduced slope-native slope). Negative values mean higher native slopes while positive values imply higher introduced slopes. Bold pvalue highlights significant differences (pv<0.05).

| **Variable** | **Slope difference** | **Pvalue** |
|---|---|---|
| **Succession time** | -0.007 | **<0.0001** |
| **Minimum colonisation time** | -0.005 | **<0.0001** |
| **Phylogenetic distance** | -0.002 | **<0.0001** |
| **Functional multivariate dissimilarity** | 0.006 | 0.364 |
| **N percentage** | 3.747 | **<0.0001** |
| **Org matter percentage** | 0.018 | **0.003** |
| **Light penetration** | 0.110 | **<0.0001** |
| **SLA dissimilarity** | -0.009 | **<0.0001** |
| **LDMC dissimilarity** | -0.002 | **<0.0001** |
| **Height dissimilarity** | -0.002 | **<0.0001** |
| **Seed mass dissimilarity** | 29.640 | **<0.0001** |
| **Burning treatment** | -0.194 | 0.285 |

**Table S8.** Differences in slope at site spatial scale between introduced and native species slopes for each variable (introduced slope-native slope). Negative values mean higher native slopes while positive values imply higher introduced slopes. Bold pvalue highlights significant differences (pv<0.05).

| Variable | Slope difference | pvalue |
|---|---|---|
| **Succession time** | -0.009 | **<0.0001** |
| **Minimum colonisation time** | -0.005 | **0.001** |
| **Phylogenetic distance** | -0.007 | **<0.0001** |
| **Functional multivariate dissimilarity** | 0.024 | 0.661 |
| **N percentage** | 0.361 | 0.858 |
| **Org matter percentage** | 0.099 | **<0.0001** |
| **Light penetration** | -0.529 | **0.032** |
| **SLA dissimilarity** | 0.015 | **0.021** |
| **LDMC dissimilarity** | -0.001 | 0.315 |
| **Height dissimilarity** | -0.013 | **<0.0001** |
| **Seed mass dissimilarity** | 51.017 | **<0.0001** |
| **Burning treatment** | 0.107 | 0.957 |

242
